# Pattern- based Contractility Screening (PaCS), a reference-free traction force microscopy methodology, reveals contractile differences in breast cancer cells

**DOI:** 10.1101/2020.05.14.097006

**Authors:** Ajinkya Ghagre, Ali Amini, Luv Kishore Srivastava, Pouria Tirgar, Adele Khavari, Newsha Koushki, Allen Ehrlicher

**Affiliations:** Department of Bioengineering, McGill University, Montreal, H3A 0E9

**Keywords:** Strain Energy, Contractility, Traction Force Microscopy, Micropatterning, Cancer metastasis

## Abstract

The sensing and generation of cellular forces are essential aspects of life. Traction Force Microscopy (TFM) has emerged as a standard broadly applicable methodology to measure cell contractility and its role in cell behavior. While TFM platforms have enabled diverse discoveries, their implementation remains limited in part due to various constraints, such as time-consuming substrate fabrication techniques, the need to detach cells to measure null force images, followed by complex imaging and analysis, and the unavailability of cells for post-processing. Here we introduce a reference-free technique to measure cell contractile work in real-time, with basic substrate fabrication methodologies, simple imaging, and analysis with the availability of the cells for post-processing. In this technique, we confine the cells on fluorescent adhesive protein micropatterns of a known area on compliant silicone substrates and use the cell deformed pattern area to calculate cell contractile work. We validated this approach by comparing this Pattern-based Contractility Screening (PaCS) to conventional bead-displacement TFM and show quantitative agreement between the methodologies. Using this platform, we measure the contractile work of highly metastatic MDA-MB-231 breast cancer cells is significantly higher than non-invasive MCF-7 cells. PaCS enables the broader implementation of contractile work measurements in diverse quantitative biology and biomedical applications.

## Introduction

Cells are not purely biochemical entities but are also subjected to physical forces and mechanics. Force-generation, sensing, and mechanical adaptation can be seen in nearly every aspect of our physiology^1,2^. Correct recognition of responses to mechanical cues are key to health, whereas dysfunctional responses are symptomatic and perhaps causative to numerous pathologies^3–6^. This signifies an urgent and pressing need to quantify how cells detect and respond to mechanical forces. This is a critical question in biology and biophysics, enabling new approaches in diagnosing and treating diverse aspects of human health.

Cell contractile forces are largely generated by molecular motors such as myosin, which pull the filamentous actin network to perform mechanical work on the surrounding matrix^7^. There are diverse methodologies to measure cell contractile work^8-10^; however, Traction Force Microscopy (TFM) has emerged as the leading approach^8-10^. TFM has revealed the roles of cell contractile forces in regulating diverse physiological and pathological processes such as cell proliferation^11^, differentiation^13^, migration^11,12^, nuclear polarization and deformation^15,16^, in virtually all adherent cells, thus making contractile work measurements a critical aspect of quantifying biological behaviors^11-16^ and potentially identifying pathologies^14^.

Although TFM is a widely useful technique, its implementation is limited in part due to experimental complexity^8-10^. TFM often utilizes protein functionalized elastic substrates, usually silicone or polyacrylamide, containing sub-micron fluorescent beads acting as fiduciary markers to capture cell-induced material deformations^8^. A typical TFM experiment involves imaging of the beads in the stressed state, followed by the detachment of cells to image the beads again to determine their positions in the unstressed state. The resulting two images are analyzed to calculate the total contractile work done by the cells in deforming the underlying substrate. Although TFM provides high-resolution traction force measurements due to the use of densely packed fluorescent beads, a limitation to this technique is lack of control over the placement and spacing of fiduciary markers^9^. These random bead arrangements necessitate the acquisition of a “null force” reference image to calculate the strain from cell-induced bead displacements, thus requiring cell detachment at the same position using enzymatic or chemical approaches^9^. This markedly complicates experimental procedures, imaging, and analysis, and precludes cellular post-processing, such as the immunofluorescence staining. While pillar-based approaches do not require a null-force image, they introduce topographical features that limit cell adhesion only to the pillar surface, thus affecting cell morphology, focal adhesions, cytoskeletal contractility, translocation of mechanosensitive proteins and stem cell differentiation^23,24^. Hence, there is a current void in cell contractility measurement methodologies that are readily implementable without affecting the biological features of the cells, thus limiting the incorporation of cell biophysics in modern quantitative biology studies.

Here, we introduce a novel reference-free approach to quantify cell contractile work based on the deformations of micropatterns of cell adhesion proteins. We print fibronectin micropatterns of known area and shape on compliant elastic silicone substrates; by imaging the cell-induced pattern area deformations and knowing substrate modulus, we calculate the total contractile work done by the cells to deform the patterns. The known pattern area makes it a reference-free approach, without introducing any specialized fabrication procedures or analysis. Continuous capture of pattern deformations allows real-time contractile measurements for longer periods of time with the ability to analyze a relatively large cell population without removing the cells, thus allowing cell post-processing of the same set of cells.

## Materials and Methods

### Synthesis of compliant silicone substrates

To measure contractile work, polydimethylsiloxane (PDMS) substrates with different stiffnesses were prepared as described previously^17,18,31^. In brief, PDMS solutions were supplied by mixing same weight ratio of component A and B of commercial PDMS (NuSil® 8100, NuSil Silicone Technologies) with different concentrations of Sylgard 184 PDMS crosslinking agent (Dimethyl, methylhydrogen siloxane, which contains methyl terminated silicon hydride units) to obtain substrates with various stiffnesses. We measured the mechanical properties of the PDMS at different crosslinker concentrations using a parallel plate rheometer (Anton Paar) and calculated the Young’s moduli (Table 1)^17,18,31^. For our experiments, 50 μl of uncured PDMS was applied to the clean 22*22 mm (No.1) glass coverslips and cured at 100°C for two hours. For traction force microscopy, prepared PDMS substrates were coated with a layer of fiduciary particles using spin coater (WS-650 Spin Processor, Laurell Technologies) and incubated at 100 °C for an hour^17,18^.

**Table 1:**
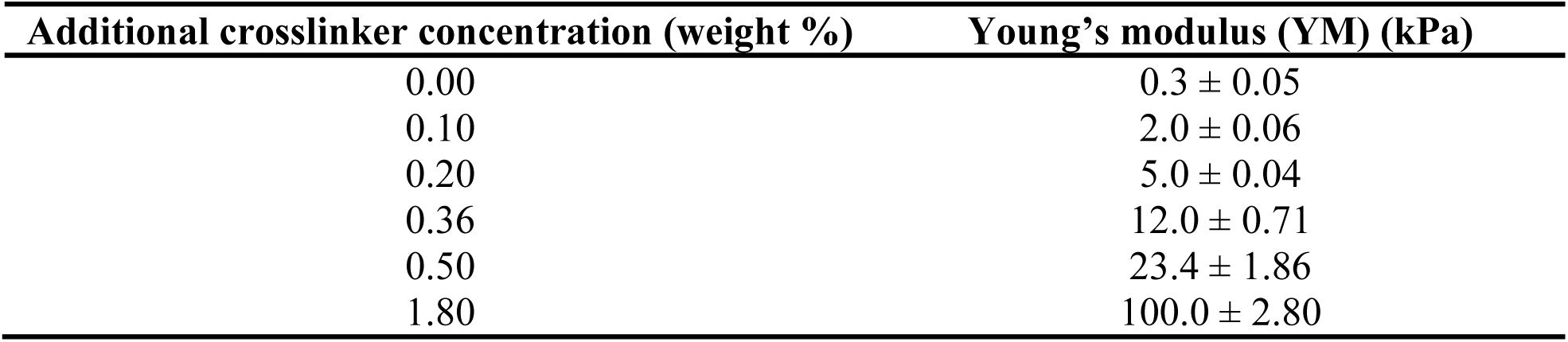
Young’s moduli for PDMS substrates containing different concentrations of Sylgard 184 crosslinking agent^17,18,31^.

### Printing on silicone substrates using UV patterning

We adhesively micropatterned silicone substrates with a UV-patterning system (PRIMO, Alveole Lab, Paris, France)^19,20^. PDMS substrates were incubated with Poly-L-Lysine (PLL, Sigma) solution (5mg/ml) prepared in 0.1M HEPES buffer (pH 8.5) for 1 hour at room temperature, followed by rinsing with MiliQ water. Positively charged PLL electrostatically adsorbs onto the negatively charged surface of silicone substrates and allows protein attachment after printing. The substrates were then incubated with Polyethylene glycol valeric acid (PEG-SVA, Laysan Bio) prepared in 0.1M HEPES buffer (pH 8.5) for 30 minutes at room temperature, followed by thorough rinsing with phosphate buffer saline (PBS) pH 7. PEG-SVA acts as an antifouling brush layer that repels protein attachment. The substrates were then covered with the UV sensitive photo-initiator solution of PLPP (Alveole Lab, Paris, France) and placed on the stage of a microscope (Nikon Ti2 Eclipse) equipped with the UV-patterning system.

To generate the patterns, we used open-source graphics software programs, Inkscape and ImageJ, to generate binary 8-bit mask image files that were loaded into PRIMO’s control software. The desired pattern was generated by a digital micromirror array in the PRIMO system and projected using a 375 nm UV laser with an intensity of 29mW/mm^2^ via 20X/0.45NA objective. The projected pattern results in localized photodegradation of the antifouling PEG-SVA brush, in the shape of the desired pattern. An exposure dose of 20 seconds was adequate to complete photodegradation of the PEG-SVA brush.

Following UV exposure, we washed the substrates with PBS and incubated them for 1 hr at room temperature with a mixture of fluorescently labeled bovine serum albumin (BSA, Alexa Fluor™ 555 conjugate, Thermofisher)(5µg/ml) and fibronectin (40µg/ml, Sigma) in PBS to adsorb the protein to the exposed PLL surface. Excess protein was rinsed off with PBS prior to cell seeding.

### Cell culture and seeding

Cell lines used in this research: NIH-3T3 (ATCC-CRL1658) mouse fibroblast cells, MDA-MB-231 (ATCC-HTB-26) highly metastatic breast adenocarcinoma cells, MCF-7 (ATCC-HTB-22) low metastatic breast adenocarcinoma cells. All cell lines were cultured in Dulbecco’s modified Eagle medium (DMEM) (Wisent) supplemented with 10% fetal bovine serum (FBS) (Wisent) and 1% Penicillin-Streptomycin antibiotic (P/S) (Thermo Fisher). Cells were seeded on the patterns for 1hour at 37°C in 5% CO_2_ environment, followed by a gentle wash with PBS to remove nonattached cells to avoid nonspecific attachments. Cells were further incubated for 16-18 hours (on patterns) before imaging at 37°C in 5% CO_2_ environment.

### Imaging

After 16-18 hours of cell seeding, cells were stained with cell tracker green CMFDA (Thermofisher) to detect cell boundaries, and the plates were transferred to a lab-built heated stage perfused with 5% CO_2_ and mounted on a confocal microscope (Leica TCS SP8 with a 10x 0.4 NA objective). With this setup, cells were imaged with transmission and fluorescence microscopy for extended periods, while maintaining a controlled culture environment.

### Immunofluorescence staining

For post-processing after contractile work measurements, we fixed the cells with 4% paraformaldehyde for 15 min at room temperature and washed three times with PBS. The cells were permeabilized with 0.1% Triton X-100 diluted in PBS for 10 minutes. To avoid any nonspecific hydrophobic binding, 2% bovine serum albumin (BSA) was added to the cells and incubated for 30 minutes at room temperature. After washing with PBS, we stained actin filaments with 10µg/ml Phalloidin (Alexa Fluor 647, Thermofisher) for 1 hour at room temperature and nuclei with 1.5 µl/ml bisBenzimide H 33342 trihydrochloride (Sigma) for 10 minutes, after which cells were washed with PBS. Fluorescence images were acquired with a Leica SP8 confocal microscope with 63X/1.4 NA oil immersion objective.

### Quantification of cell contractile work

To measure cell contractile work, we applied the following equation to calculate strain energy from pattern area deformations and material properties of the silicone substrates:

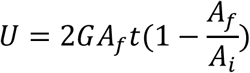

Where U, *A*_*i*_, *A*_*f*_, G and *t* are total strain energy, initial pattern area, deformed pattern area, substrate shear modulus and substrate thickness respectively (derivation in Supporting Material). In brief, we measure the deformed and undeformed pattern area by thresholding the fluorescent pattern images. For strain energy calculations, we use a single averaged value of undeformed pattern area (2401.96 ± 32.24 µm^2^, Fig S1a) and compare it with the cell deformed pattern area, which along with known modulus of silicone substrate allows us to calculate total strain energy applied by the cells to deform the underlying substrate. We calculate outlier strain energy values from undeformed pattern areas on each stiffness substrates (Fig S1b).

### Data analysis

Cell strain energy was calculated using a custom MATLAB script which requires fluorescent pattern images, substrate stiffness and initial pattern area. The code calculates the pattern area and strain energy values for the respective cells. The code is available on the GitHub repository with experimental details and example data for analysis (https://github.com/ajinkyaghagre/PaCS_matlabcode). Further relevant data are available from corresponding authors request.

## Results

### PaCS Design and Analysis

In PaCS, adhesive protein micropatterns are printed on the surface of compliant silicone substrates. We chose polydimethylsiloxane (PDMS) because of its favorable material properties such as the ability to tune its stiffness over a large physiological range, chemical stability (nondegradable) and bioinertness^17,18^. PDMS is also optically transparent (refractive index ∼1.4) and amenable to spin coating. This facilitates creating a uniform and flat surface that avoids the confounding effects of hydrogel porosity on the cells^17,18^.

We cured flat PDMS substrates of specific Young’s moduli (E = 2 ± 0.06, 12 ± 0.71 and 23.45 ± 1.86 kPa) on cover glass as described previously^17,18^. Next, we printed adhesive protein micropatterns of desired shapes and sizes on PDMS substrates using the PRIMO photopatterning system^19,20^.

In brief, the PDMS substrates were first coated with PLL which promotes cell attachment, followed by a coating with an antifouling agent PEG-SVA. The PRIMO photopatterning system utilizes a UV laser which projects the desired pattern on the surface of the PDMS substrates. The projected UV laser etches PEG-SVA in the presence of a photo initiator PLPP, thus exposing the underlying PLL layer for adhesive protein attachment (Fig 1a). Using this system, we confined NIH 3T3 fibroblast cells on square micropatterns of ∼2400µm^2^ printed on 2 kPa PDMS substrates (Fig 1b). We used a combination of fluorescent BSA and fibronectin for pattern visualization and cell attachment. Fibroblast cells deform the soft silicone substrate using contractile forces thus deforming the printed patterns into arbitrary shapes (Fig 1b). We binarize the images to measure the deformed pattern area and compare it with the initial pattern area (Fig 1c), which along with material properties of silicone allows us to calculate total contractile work done (strain energy) by the cell to deform the substrate (Fig 1d, Supporting Material). In this case, the representative fibroblast cell is applying a strain energy of 0.132 ± 0.01 pJ, deforming the square pattern of 2401.96 ± 32.24 µm^2^ to a pattern area of 2003.68 ± 44.31µm^2^, on a 2 kPa PDMS substrate (Fig 1b, c).

**Figure 1:**
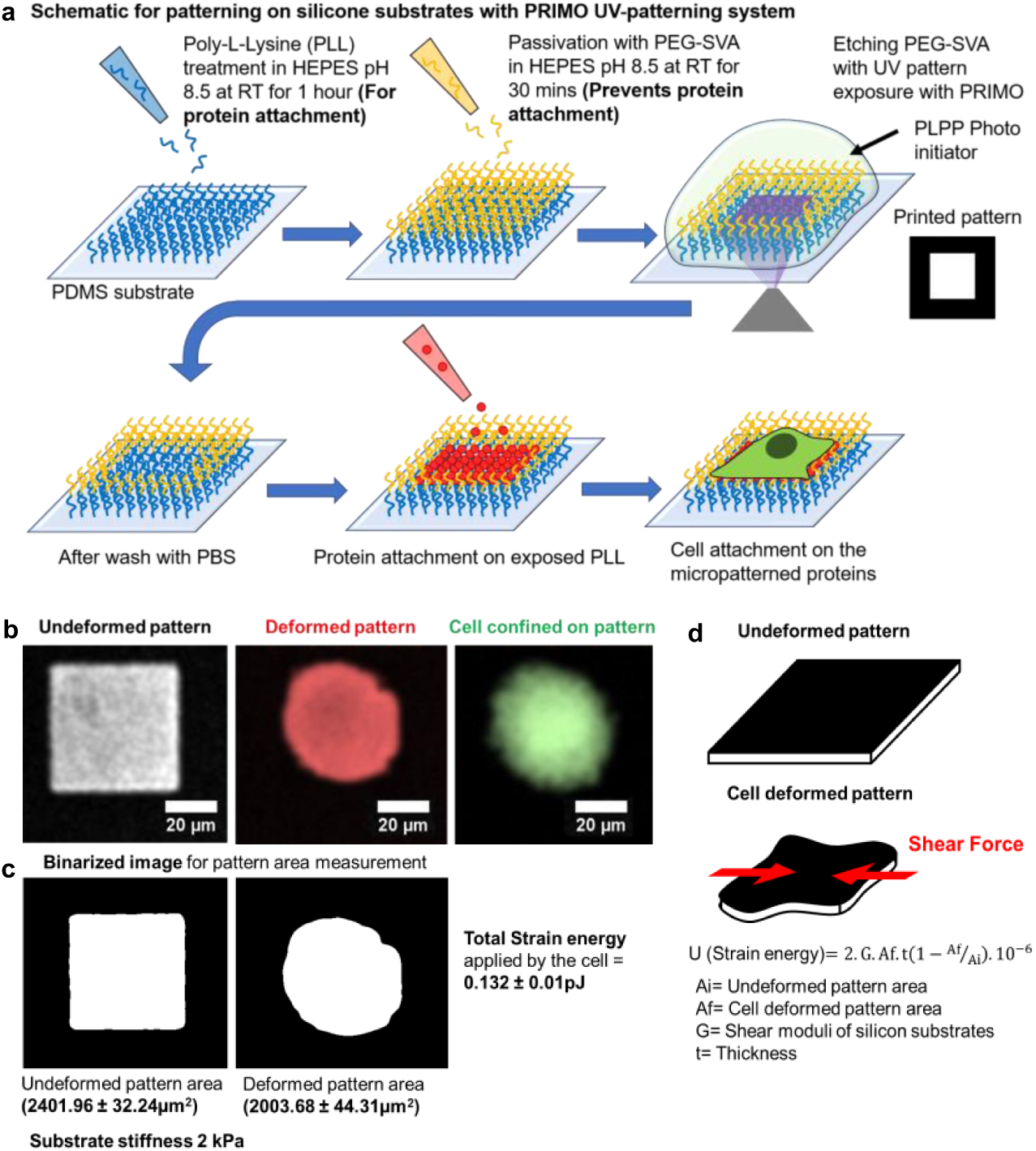
Cells deform adhesive protein patterns on compliant silicone substrates: **a**, Schematic for patterning on silicone substrates with PRIMO photopatterning system. **b**, Fluorescent BSA and fibronectin square pattern on 2 kPa PDMS substrates undeformed (white) and deformed by the cell (red). Cell on the pattern stained with CMFDA cell tracker (green). **c**, Binarized image of the undeformed and deformed pattern used for area measurements, used to calculate total strain energy applied by the cells to deform the underlying pattern, in this case 0.132 ± 0.01 pJ. **d**, Schematic explanation for strain energy calculations using the volumetric strain approach.

### PaCS accurately captures contractile work across diverse cell shapes

To determine the accuracy of our contractile work measurement, we measured cell strain energy using both conventional bead-based TFM and PaCS simultaneously. We coated the PDMS substrates with fluorescent beads, followed with printing of adhesive micropatterns on the substrates. Fluorescent beads allow us to measure contractile work with bead-displacement TFM and compare it with the contractile work calculated from PaCS for the same cells. We printed micropatterns of the same area (∼2400µm^2^) but with different shapes (square, circle, triangle, rectangle, star, and pentagon) on PDMS substrates with a Young’s modulus of 12 kPa and measured cell contractile work of NIH 3T3 fibroblast cells with both bead-displacement TFM and PaCS.

Fibroblast cells confined on diverse pattern shapes deformed the patterns, the areas of which were used to quantify cell contractile work (Fig 2a, S2a). When compared with bead-based TFM measurements, PaCS accurately measured contractile work of fibroblast cells confined on patterns of all investigated different shapes (Fig 2b). The increased pattern deformations strongly correlate with higher strain energies applied by the cells to deform the underlying substrate (Fig S2b). Thus, PaCS accurately and precisely measures cell contractile work irrespective of pattern shape with the availability of the cells for post processing.

**Figure 2:**
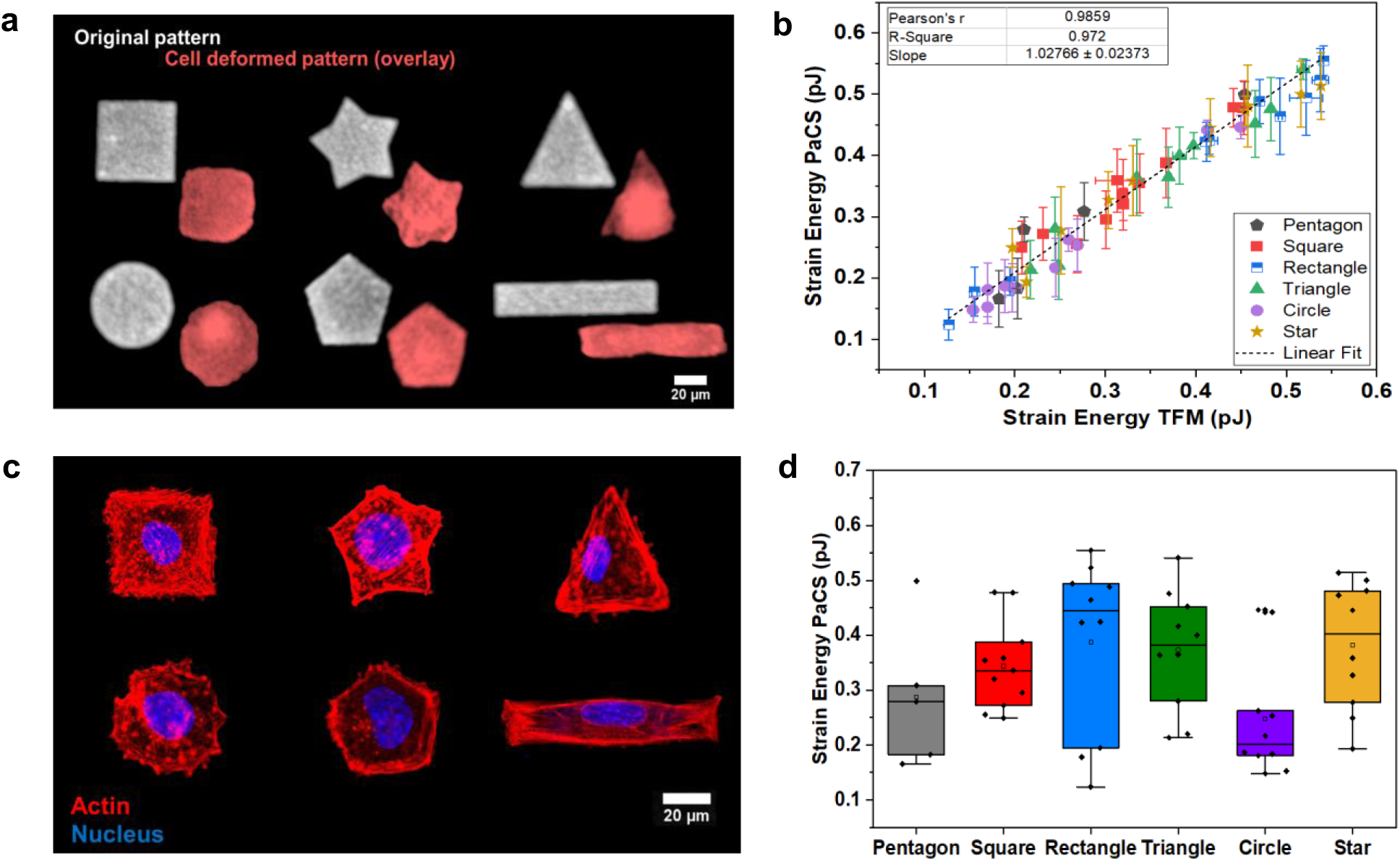
PaCS accurately measures cell contractile work across diverse cell shapes: **a**, Fluorescent BSA-fibronectin patterns of various shapes deformed by 3T3 Fibroblast cells on 12 kPa PDMS substrates. **b**, Total strain energy calculated with PaCS strongly correlates with strain energy calculated with TFM (n=56). **c**, 3T3 Fibroblast cells fixed and stained with phalloidin (actin) and DAPI (nuclei), imaged on Leica SP8 confocal (63X/1.4NA objective). **d**, PaCS strain energy for 3T3 Fibroblast cells indicate cells confined in circular shape to apply the least strain energy.

To demonstrate the ability of this technique to allow post-processing, we fixed and stained the confined cells with phalloidin and DAPI to visualize the actin filaments and nucleus, respectively (Fig 2c). Consistent with previous work^27-30^, these fluorescent images reveal that actin filaments are most concentrated on external polygon edges and terminate at polygon vertices. Conversely, circular shapes appear to promote radially aligned actin filaments in the cell, which results in cell applying lower strain energies when compared with other cell shapes (Fig 2d). These data show how profoundly cell geometry impacts cytoskeletal structure.

### PaCS resolves time-dependent contractile work and cytoskeletal activity as a function of substrate stiffness

To examine the accuracy of PaCS on different moduli substrates, as before we compared PaCS measurements with bead-based TFM using PDMS substrates coated with fluorescent beads. We measured the contractile work of 3T3 fibroblast cells on square patterns of the same area (∼2400µm^2^) printed on PDMS substrates with Young’s moduli of 2, 12 and 23.45 kPa. We observed decreased pattern deformation with increasing substrate stiffness (Fig 3a, S3b), and that cells applied more contractile work on stiffer substrates (Fig 3b). The contractile work measurements from PaCS are highly correlated with the work calculated with TFM (Fig 3b). Thus, PaCS accurately and precisely resolved contractile differences of cells on different stiffness PDMS substrates (Fig S3c).

**Figure 3:**
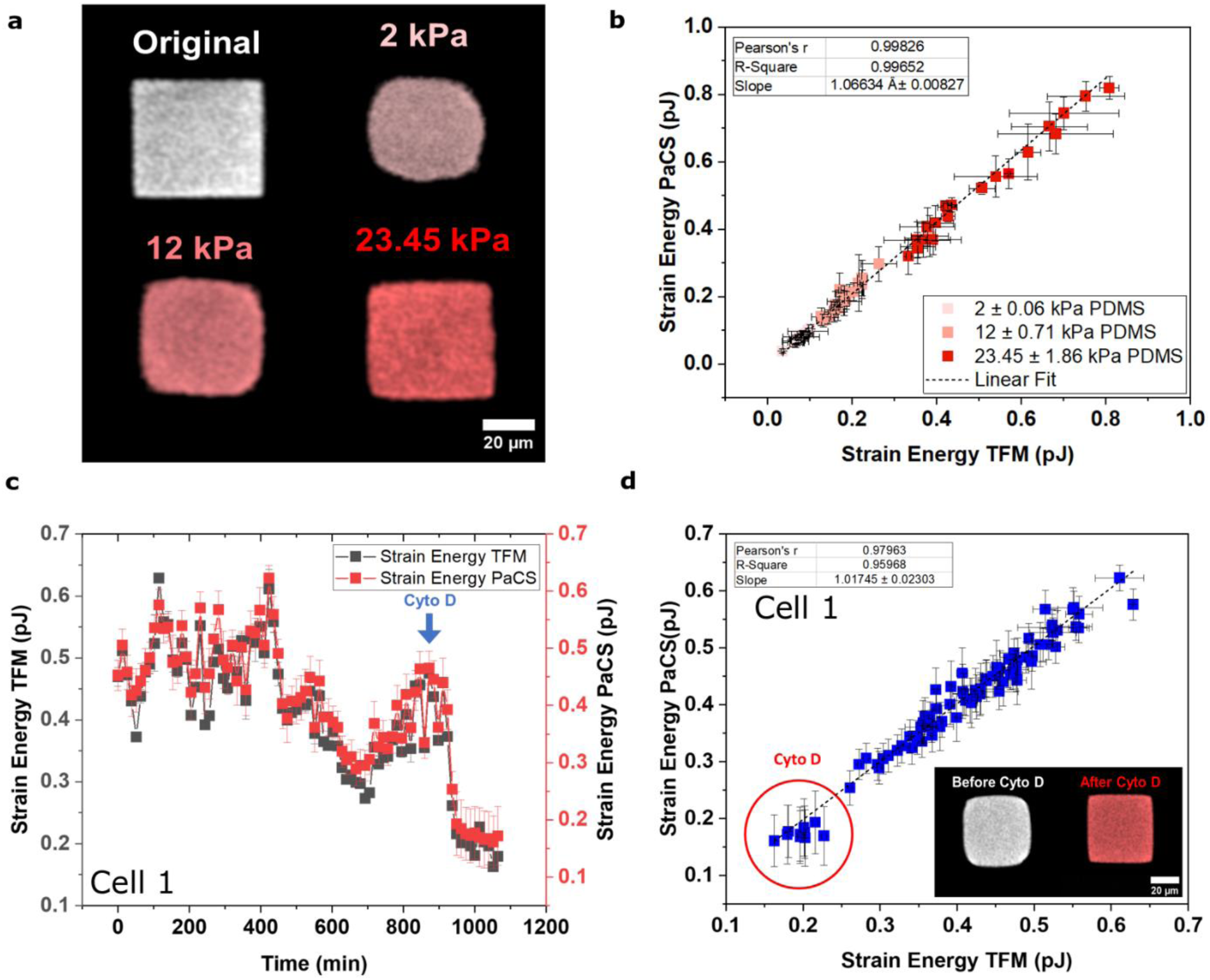
PaCS resolves time dependent contractile work and cytoskeletal activity as a function of substrate stiffness. **a**, Pattern deformations by 3T3 fibroblast cells on PDMS substrates of different Young’s moduli (2, 12 and 23.45 kPa) **b**, Total strain energy calculated with pattern deformations strongly correlates with strain energy calculated with TFM (n=60). **c**, Time-dependent relation between strain energy calculated with PaCS and TFM (cell 1), blue arrow represents the time of addition of Cyto D (Time interval 12.83mins) (n=84 timepoints). **d**, Strain energy calculated with pattern deformations with time strongly correlates with strain energy calculated with TFM, red circle represents the strain energy values after Cyto D (n=84 timepoints).

**Figure 4:**
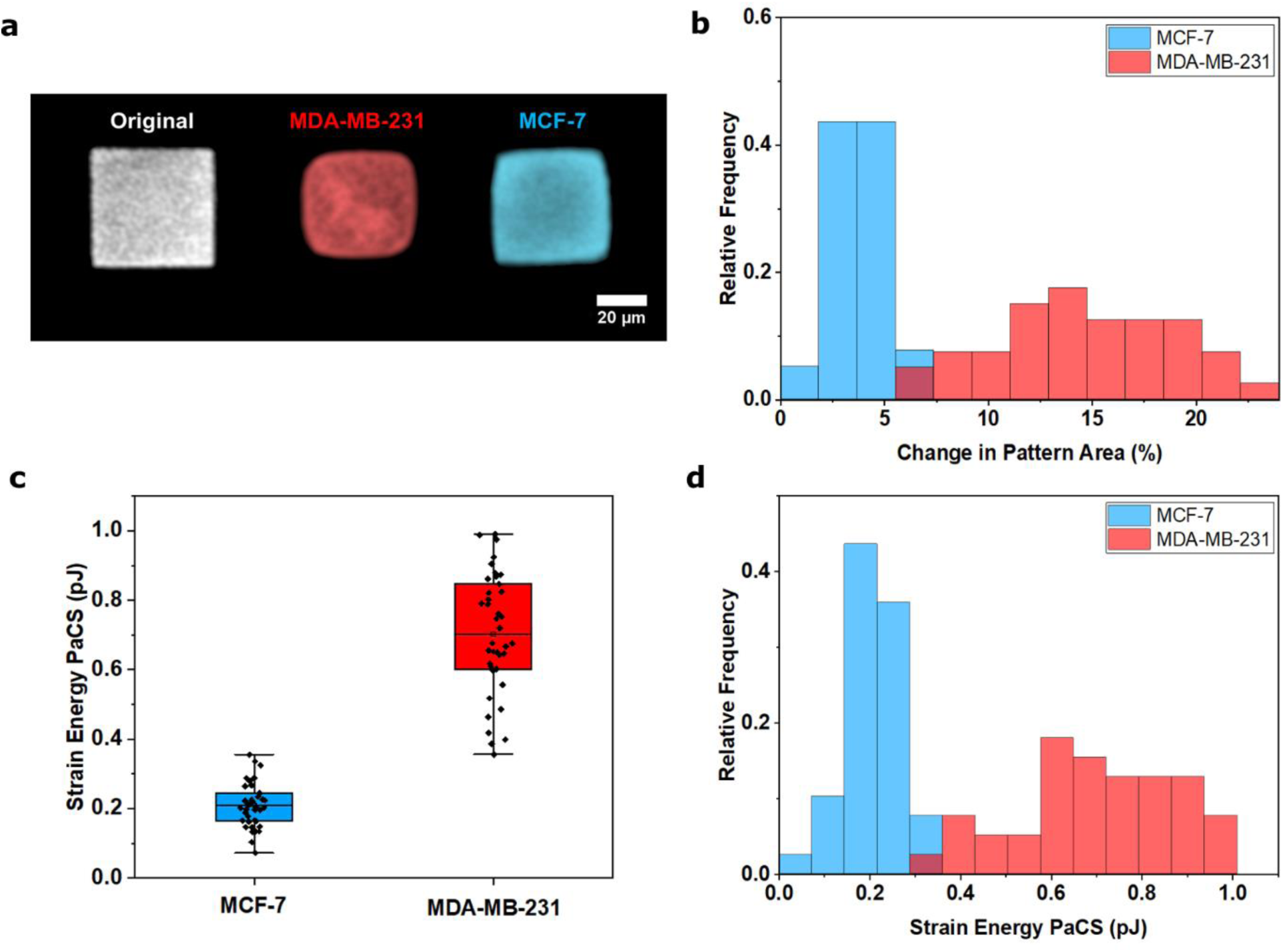
PaCS reveals contractile differences between low metastatic and highly metastatic breast cancer cells. **a**, Pattern deformations for high metastatic MDA-MB-231 (red) and low metastatic MCF-7 (blue) cells, show higher deformations by highly metastatic cells. **b**, Histogram of percent change in pattern area for MDA-MB-231 (n=40) and MCF-7 cells (n=39). **c**, PaCS Strain Energy for MDA-MB-231 cells (n=40) and MCF-7 cells (n=39). **d**, Histogram of strain energy for MDA-MB-231 cells (n=40) and MCF-7 cells (n=39) reveal contractile differences between the cell lines, with highly metastatic cell being more contractile.

We further tested the ability of PaCS to resolve time-dependent contractile work. We measured fibroblast cell contractile work on square patterns (∼2400µm^2^) on PDMS substrates (12 kPa) for 18 hours; to further test the sensitivity of this technique we inhibited the cell contractile work by depolymerizing actin using Cytochalasin D (Cyto D) in the last few hours of measurement. Using time-dependent PaCS, we accurately measured cell contractile work with time when compared to TFM measurements, even after the Cyto D treatment (Fig 3c and d, S4). The increased pattern area after drug treatment, indicates decreased contractile work of the cell with time. (Fig 3d, inset,S4)

Hence, PaCS accurately and precisely measures time-dependent contractile work as a function of substrate stiffness, without the need for a null force image, thereby enabling higher number of measurements per experiment, with the availability of the cells for post-processing such as immunofluorescence.

### PaCS reveals contractile work differences between metastatic breast cancer cells

To exemplify potential applications of this technique, we performed contractile work measurements using PaCS on low and highly metastatic breast cancer cells. The tumor microenvironment undergoes diverse mechanical and chemical changes throughout the neoplastic progression^21,22^. During cancer metastasis, cells form the primary tumor site acquire the ability to escape and migrate through the heterogeneous tumor microenvironment to establish secondary tumors. Despite being linked to poor prognosis, there are few direct biophysical clinical tests available to diagnose the likelihood of metastasis^21,22^. Because metastasis of most solid tumors requires cells to exert force to reorganize and navigate through the dense stroma, and has been previously correlated with contractility^21,22^, we investigated the differences in cellular force generation between low and highly metastatic cancer cells with PaCS. In this study we measured contractile work of highly metastatic (MDA-MB231) and weakly metastatic (MCF-7) breast cancer cells using square patterns (∼2400µm^2^) printed on 12 kPa PDMS substrates. Highly metastatic MDA-MB-231 cells exhibited higher pattern deformation than low metastatic MCF-7 cells (Fig 5a, b and S5). In agreement with previous findings, we observed that highly metastatic cancer cells exerted larger strain energies than breast cancer cells with lower metastatic potential (Fig 5c, d)^21^. These results demonstrate the ability of PaCS to resolve contractile changes between different cancer cell types, which may lead to simplified biophysical clinical tests to diagnose cancer metastasis.

## Discussion

PaCS combines the approaches of adhesive protein micropatterning on soft silicone materials with automated image analysis to provide real-time cell contractile work measurements. The soft silicone base offers tunable stiffness in physiological range^17,18^, along with controlled cell confinement using adhesive protein micropatterns, broadens its applications across multiple cell types and functions.

We demonstrate the ability of PaCS to measure cell contractile work of fibroblast cells across diverse shapes and highlight its potential to resolve increasing contractile work of cells on increasing substrate stiffness. The correlation with conventional TFM revealed high accuracy of PaCS in measuring contractile work across all diverse shapes and substrate stiffness.

Although conventional TFM and micropillar techniques provide multi-dimensional force resolutions in the form of vectors assigned to specific focal adhesions, such techniques are limited to specific biophysical questions for a limited number of cells^8-10,26,27^. Such techniques require high resolution imaging to resolve small scale forces and demands extensive workflow,which comes at a cost of the limited number of measurements and simplicity. PaCS is a reference-free platform, that measures contractile work from a single image of deformed patterns, thus simplifying the experimental workflow and increasing the number of measurements per experiment.

The use of unconfined cells in TFM and micropillar techniques further complicates the workflow with the requirement of precise position monitoring of the cells moving within each time frame. In PaCS cells are restricted to a single position and do not migrate which simplifies further imaging and analysis. Time-dependent PaCS measurements of fibroblast cells on silicone substrates, revealed a strong correlation of contractile work with TFM, and highlighted the time-dependent sensitivity of the technique after treatment with contractile inhibitor.

Moreover, micropillar and microdot techniques complicate the measurements by introducing additional topographical features on the substrate surface, along with variations in distance between the dots or pillars, all of which have shown to affect the biology of the cells^23,24^. In PaCS cells are on confined adhesive patterns on continuous substrates, thus avoiding any effect from topographical features of the substrate.

We further demonstrate a potential application of PaCS in resolving contractile work differences in cancer cells. During cancer metastasis, the neoplastic microenvironment not only confers biochemical changes but also alters the biophysical phenotype of cancer cells^25^. Malignant cells are reported to be highly contractile with increased migration and compliance. Such biophysical markers can be used to diagnose metastatic potential of cancer cells or can be screened to develop treatments that directly target these biophysical characteristics and thus effectively hinder metastasis. Contractile forces are emerging as biophysical markers in the majority of cancer models including breast, prostate, lung and bone^21,25^.

In this study we used PaCS to measure contractile work differences between highly metastatic (MDA-MB-231) and weakly metastatic (MCF-7) breast cancer cells. In agreement with the previous finding, PaCS detects high contractile work done by invasive MDA-MB-231 breast cancer cells, when compared to less invasive MCF-7 cells, thus highlighting the involvement of contractility in cancer metastatic progression. The ability of PaCS to detect contractile work differences across multiple cell types, broadens the horizon of its applications.

Recently contractile forces have been implemented as a crucial parameter for anti-cancer and bronchoconstrictor drug screenings, thus demonstrating the potential of PaCS for drug screening applications^22,26^. With further advancements PaCS has the potential to become a leading technology for the diagnosis of diseased conditions involving aberrant cellular force generation.

PaCS can be used to study the physiological role of contractile forces in regulating biological processes such as cell migration, proliferation, stem cell differentiation and nuclear deformation. Such studies need to be explored further to understand the complexity of cell biological processes.

However, such widespread use of cell mechanics across cell biology and medicine is limited by the inadaptability of current approaches across multiple labs. The standard techniques such as TFM are too difficult too be operated by non-experts, thus creating a demand for simple adaptable techniques that can be operated by anyone. PaCS provides a quick and accurate contractile work measurement with a simple workflow, which makes it readily adaptable across multiple labs. While we have demonstrated this approach using UV micropatterning, we anticipate that broader implementation could also be achieved with soft-lithography based microcontact printing^32^.

## Conclusion

In this paper we introduce a new technique to measure cell contractile work using adhesive pattern deformations. This technique allows for real-time cell contractile measurements with simpler fabrication protocols and analysis. Using this technique, we revealed contractile work differences of fibroblast cells confined on different shapes and substrate stiffness. We measured differences in contractile work with the time and observed the differences after drug-induced contractile inhibition. Finally, we demonstrate the application of this technique in resolving the contractile work differences in benign and metastatic breast cancer cells. The ability of this technique to measure contractile work in real-time from the pattern deformations fills a current void in simplified cell contractile methodologies, and provides a promising future for the incorporation of cell biophysics in broader quantitative biology studies.

## Supporting information

Supplementary Video_Cell1

## Acknowledgements

AJE acknowledges grant support from NSERC (RGPIN/05843-2014, EQPEQ/472339-2015, RTI/00348-2018), Canadian Cancer Society Grant #703930, Canadian Foundation for Innovation Project #32749, Prostate Cancer Canada D2019-2180, the Canada Research Chairs program, and CIHR # 143327. AG, AA, LKS, PT, NK were partially supported by McGill Engineering doctoral awards. The authors thank Dr Peter Seigel for the kind gifts of breast cancer cell lines MDA-MB-231 (ATCC-HTB-26) highly metastatic breast adenocarcinoma cells, MCF-7 (ATCC-HTB-22) low metastatic breast adenocarcinoma cells.

