## Supplementary Information for "Pattern- based Contractility Screening (PaCS), a reference-free traction force microscopy methodology, reveals contractile differences in breast cancer cells"

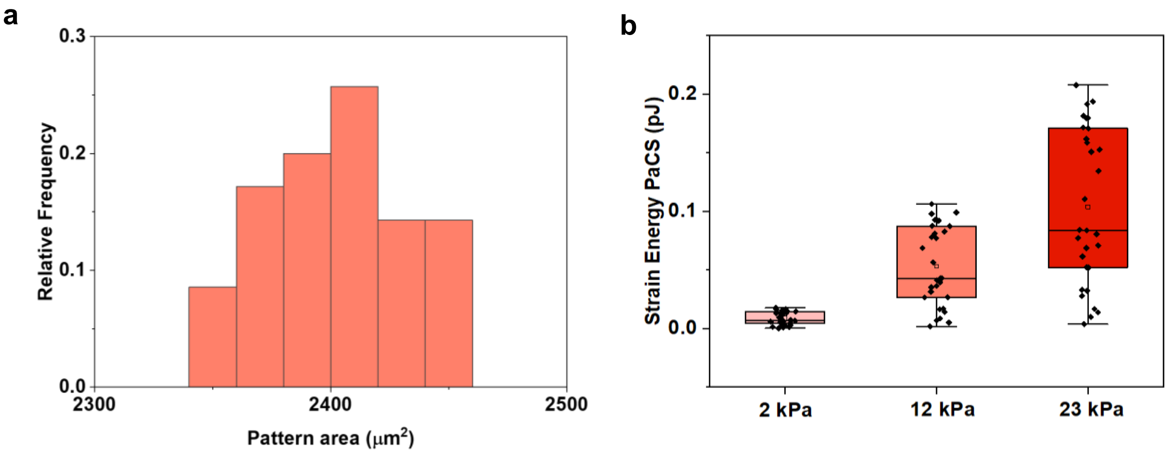


**Figure S1: a,** Area distribution of printed square patterns of 2400μm$^2$ on soft silicone substrates. **b,** Baseline strain energy values for respective substrate stiffness.

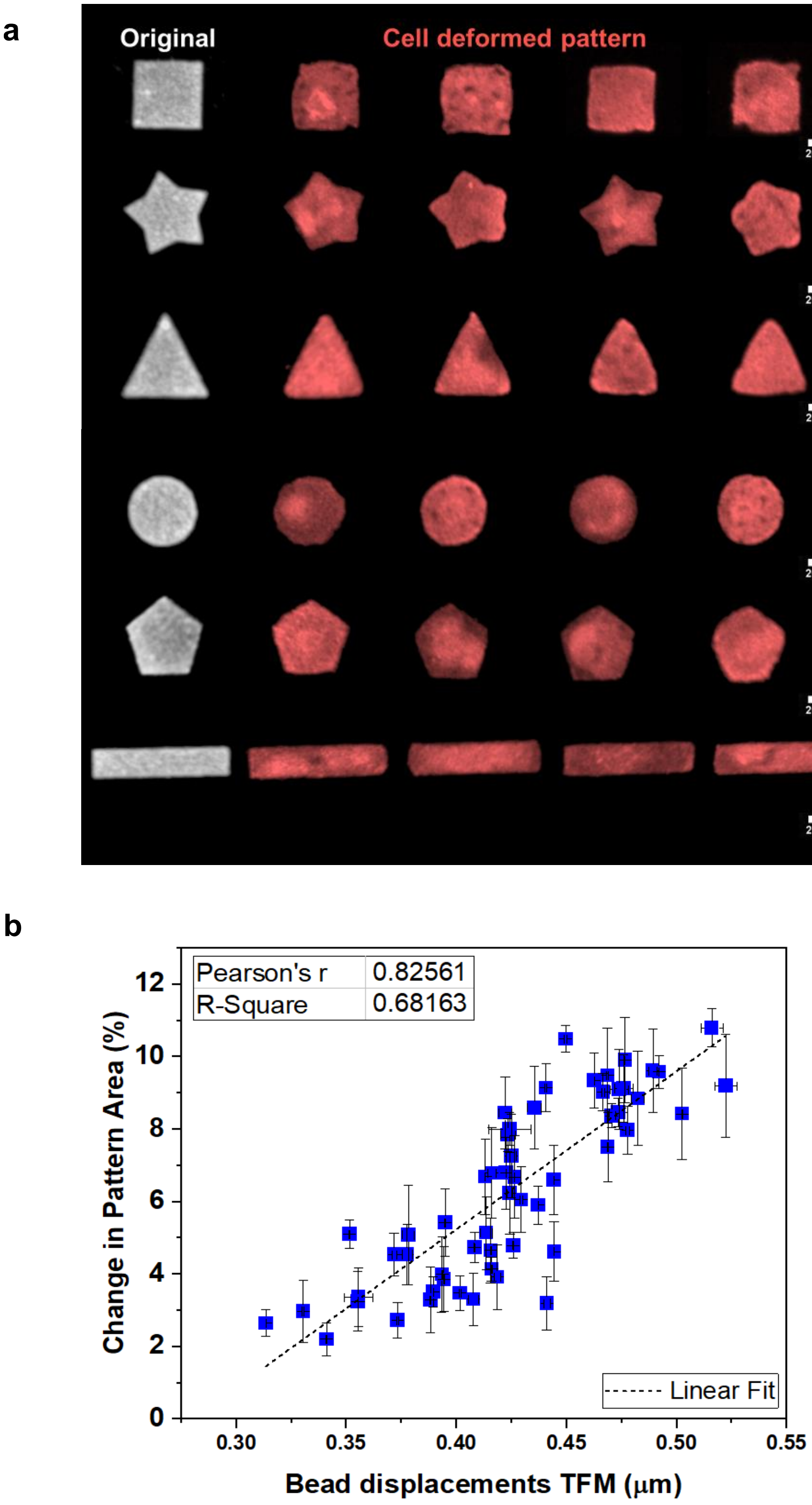


**Figure S2: a,** Pattern deformations by 3T3 Fibroblast cells for various pattern shapes. **b,** Plot representing the high correlation between strain energy and percent change in pattern areas (n=56).

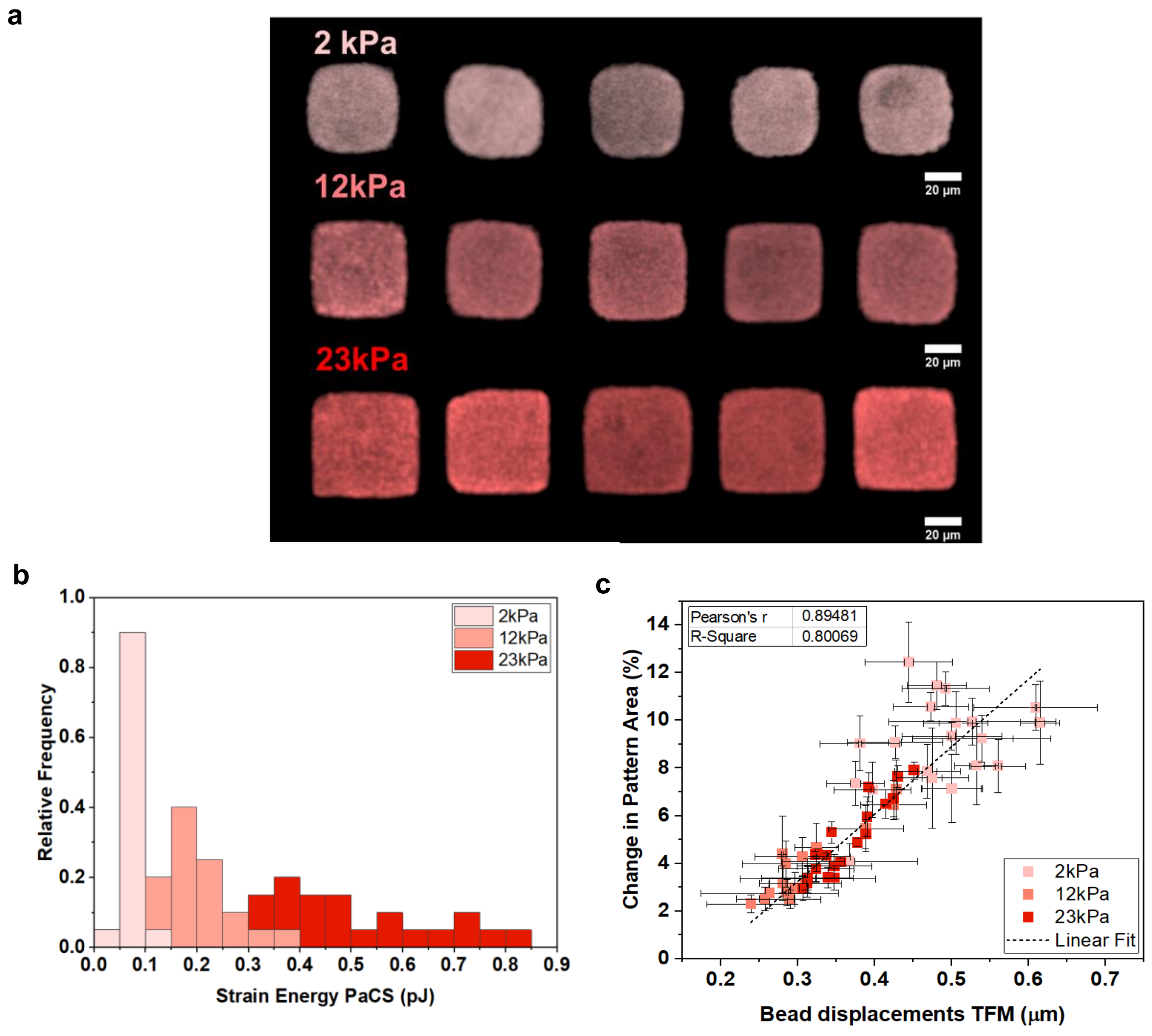


**Figure S3: a,** Pattern deformations by 3T3 Fibroblast cells confined on square patterns on different stiffness silicone substrates. **b,** Distribution of PaCS strain energy for cells on different stiffness silicone substrates. **c,** Plot for percent change in pattern area and TFM bead displacements for fibroblast cells on different stiffness silicone substrates.

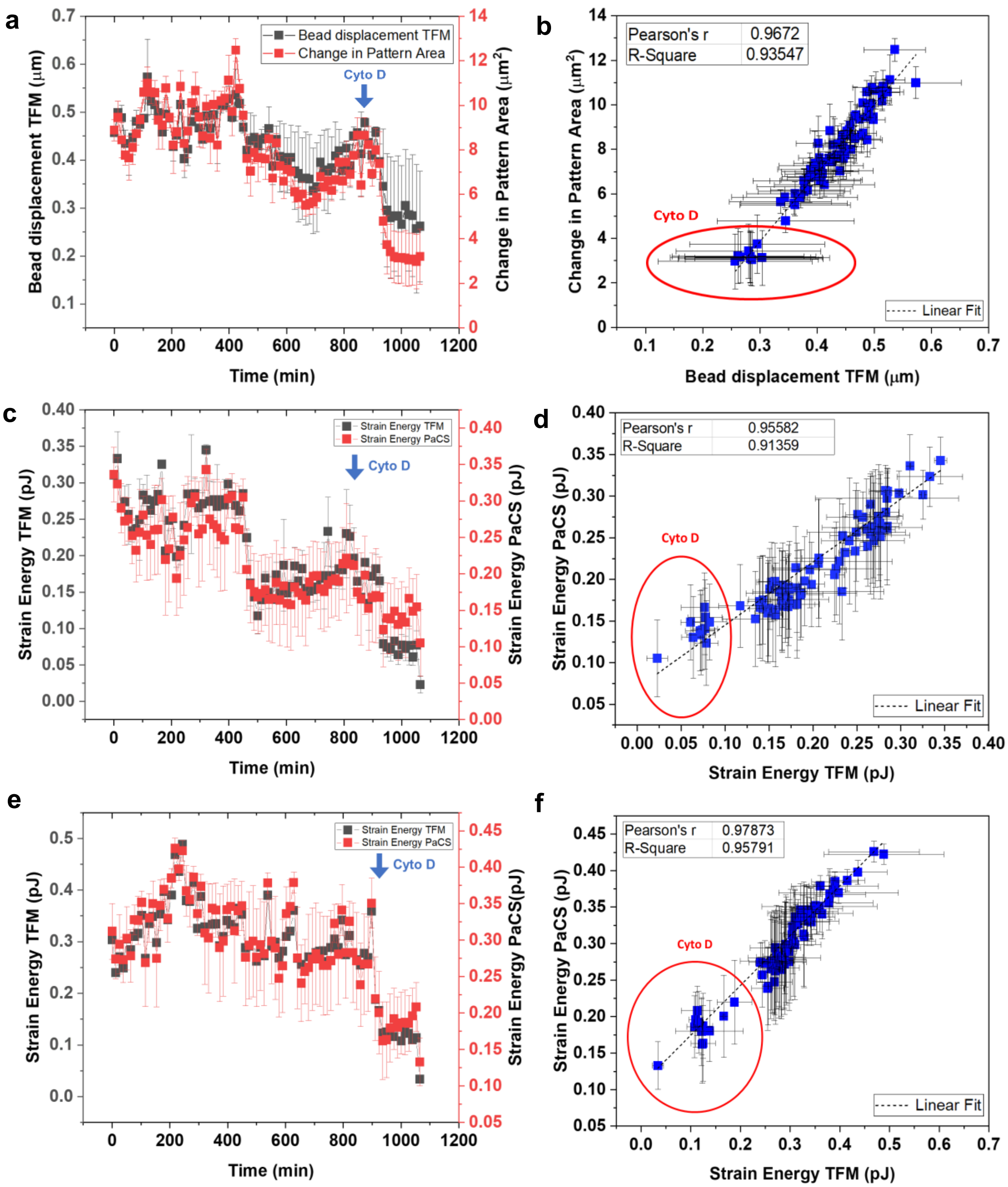


**Figure S4: a,** Time dependent relation between TFM bead displacement and percent change in pattern area with time for cell1, blue arrow represents the time of addition of Cyto D (Time interval 12.83mins) (n=84 timepoints). **b,** TFM bead displacements strongly correlate with percent change in pattern area with time, red circle represents the datapoints after addition of Cyto D (n=84 timepoints). **c and e,** Time dependent relation between strain energy calculated with PaCS and TFM (cell2 and cell 3), blue arrow represents the time of addition of Cyto D (Time interval 12.83mins) (n=84 timepoints), **d and f,** Strain energy calculated with pattern deformations with time strongly correlates with strain energy calculated with TFM (cell 2 and cell 3), red circle represents the strain energy values after Cyto D (n=84 timepoints).

a

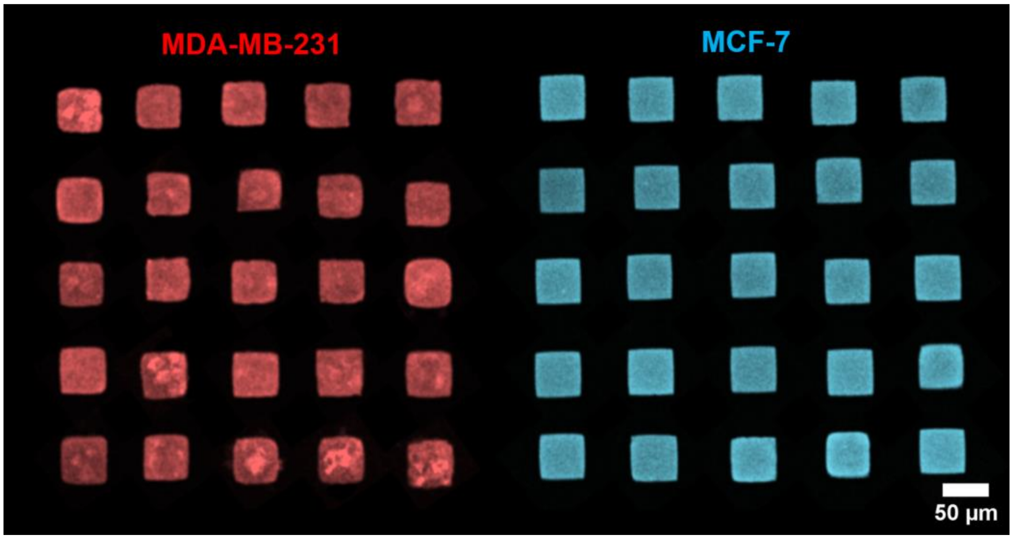


**Figure S5: a,** Pattern deformations by MDA-MB-231 and MCF-7 cells confined on square patterns on 12kPa PDMS substrates, were MDA-MB-231 cells are showing higher pattern deformations than MCF-7 cells.

**Video S1:** Time dependent pattern deformations for cell 1 captured for 18hours and treated with Cyto D in the final hours (Time interval: 12.83 mins, 84 frames)

**Appendix:**

Derivation for total strain energy using volumetric strain approach:

We consider a simple case of unidirectional pure shear on a substrate, where strain energy density is formulated as

$$u = \frac{1}{2}\tau\gamma \qquad (1)$$

Where τ and γ are shear stress and shear strain respectively. Considering the constitutive law ($\tau = G\gamma$ ), the equation 1 could be written as follow

$$u = \frac{1}{2}G\gamma^2 \qquad (2)$$

On the other hand, one can define volumetric strain in a plane as follow

$$\frac{\Delta V}{V} = \epsilon_x + \epsilon_y - {\epsilon_{xy}}^2 \qquad (3)$$

Where $\epsilon_x$ , $\epsilon_y$ , and $\epsilon_{xy}$ are normal in x direction, normal in y direction and shear strains respectively. By combining equations 2 and 3 for pure shear, and taking into consideration that $\gamma = 2\epsilon$, we have

$$\frac{\Delta V}{V} = \frac{-u}{4G} \qquad (4)$$

By assuming $\frac{\Delta V}{V} \approx \frac{\Delta A}{A}$ the total strain energy could be calculated as

$$U = 2GA_f t(1 - \frac{A_f}{A_i}) \qquad (5)$$

Where $A_f$, $A_i$, and *t* are initial area, final deformed area, and substrate thickness respectively. For the thickness we consider a value of 0.3µm on which observed an accurate measurement of strain energy when compared with TFM and is also the maximum range of z displacements for the cells on 2D substrates[10].
